# Exploring the Conformational Landscape of Adenylate Kinase and Beyond: A Benchmark of Protein Folding Models

**DOI:** 10.1101/2025.11.04.686486

**Authors:** Aryan Bhasin, Antoine Delaunay, Francesco Saccon, Yunguan Fu

## Abstract

Protein folding models have revolutionized structure prediction but struggle to capture conformational flexibility. Recent studies perturb inputs or parameters to sample alternative conformations, while diffusion-based approaches generate conformational ensembles directly. Although the former have been benchmarked to some extent, the latter have yet to be evaluated, and sub-domain dynamics validation remains limited. Here, we present a systematic benchmark of nine methods across 20 monomeric proteins with active and inactive states. We extend the pairwise aligned error metric to ensembles and reveal that protein identity exerts a non-negligible influence on model performance. Focusing on Adenylate Kinase, a well-studied enzyme with extensive molecular dynamics (MD) data, we find that Chai-1 performs the best in recovering known conformations, identifying mobile regions, and capturing transition trajectories. These results highlight the potential of generative models as efficient alternatives to MD for exploring protein conformational dynamics and provide a rigorous benchmark for dynamic structure prediction.

## 1 Introduction

Proteins carry out essential biological functions governed by their three-dimensional structures. Many proteins are not static, but instead can be found in multiple stable conformations depending on their environment and function, in which allosterically-mediated conformation changes permit advanced functions controlled by responses to different stimuli [1]. These conformational shifts are often the physical basis of a protein’s function. For instance, they drive the circadian clock through the rhythmic changes in KaiB [2], enable efficient oxygen transport by hemoglobin [3], and facilitate photoprotection in plants via the pH-induced dynamics of PsbS [4]. A well-studied example is Adenylate Kinase (AdK), which cycles between open and closed conformations to regulate cellular energy balance [5]. Substrate binding induces large-scale domain movements that transition AdK from an open, inactive form to a closed, catalytically active state. Accurately modeling these structural transitions is key to understanding how proteins perform their functions, and how those functions might be disrupted or modulated. Dysfunction in AdK has been linked to severe immunodeficiencies [6] and cancer [7], underscoring the biological importance of conformational control.

Experimental techniques such as X-ray crystallization, nuclear magnetic resonance (NMR) spectroscopy and cryo-electron microscopy (cryo-EM) can provide insights into protein structure and dynamics, but they often capture only static snapshots, making it difficult to resolve multiple conformational states. NMR is limited by resolution and throughput, while cryo-EM faces challenges with conformational heterogeneity and time resolution [8, 9]. Molecular dynamics (MD) simulations are frequently used to complement these approaches, offering detailed models of conformational transitions, but are computationally expensive [10].

Protein folding models such as AlphaFold2 [11] and AlphaFold3 [12] predict protein structures directly from sequence with unprecedented accuracy, speed, and scalability. The introduction of AlphaFold2 has been widely recognized as a paradigm shift in biological research [13]. Building on this foundation, AlphaFold3 achieves atomic-resolution modeling of complex cellular assemblies, capturing interactions among proteins, nucleic acids, and small molecules within native subcellular contexts [14]. Several open-source reproductions of AlphaFold3, including Chai [15, 16] and Boltz [17, 18], now deliver comparable levels of accuracy. Collectively, these computational advances have become indispensable in laboratories worldwide, reshaping strategies for drug discovery and molecular biology [19].

These protein folding models were often trained to return a single, confident structure representing the most likely/native-like structure for a given sequence, and thus are limited in their ability to probe functionally relevant conformational diversity. Multiple studies modified or perturbed the multiple sequence alignment (MSA) input features to sample different conformations by disrupting the co-evolutionary signals, comprising sub-sampling of the MSA [20], clustering of the MSA [21], random masking of MSA [22] and alanine-mutagenesis of random residues in the MSA [23]. Other methods used random dropout of model weights in both the Evoformer and structure modules during inference [24], or re-trained AlphaFold2 with known structural clusters of sequences [25]. Recent diffusion-based approaches, including AlphaFold3 and its open-source reproductions, were designed to be generative and can sample multiple structures from the same input.

While these diverse strategies show promise, they were typically evaluated on different protein targets using disparate metrics. Furthermore, these studies mostly focused on overall conformational changes rather than exploring sub-domains that experience conformational changes, with limited comparison to molecular dynamics. Notably, diffusion-based folding models have yet to be evaluated for their ability to sample alternative conformations. Here, we present a systematic and head-to-head benchmark, evaluating nine state-of-the-art conformational sampling methods across a curated set of 20 monomeric proteins with known active and inactive states. We focus on Adenylate Kinase (AdK) for in-depth analysis, given its well-characterized, large-scale conformational transitions and the wealth of available experimental and simulation data. We assess each method on its ability to recover known end-states, predict plausible intermediate conformations along the transition pathway, and identify the flexible “mobile” domains driving the transition, using an extension of the aligned error (AE) matrix computed across ensembles of predicted structures. Our results reveal a clear performance hierarchy, and we demonstrate that AlphaFold3-based generative models, particularly Chai-1, are uniquely capable of not only predicting both stable states with high accuracy but also capturing a transition trajectory between them. This finding positions these next-generation models as a compelling and computationally efficient alternative to traditional molecular dynamics simulations for probing the conformational landscapes that underpin protein function.

## 2 Results

### 2.1 Benchmarking nine sampling methods against twenty protein targets including enzymes, transport, and chaperone proteins

To evaluate how well state-of-the-art folding models capture conformational diversity, we benchmarked nine representative methods on a curated dataset of 20 proteins. The sampling methods included AlphaFold2, AlphaFold2-Dropout, MSA masking, MSA alanine mutation, MSA subsampling, MSA clustering, Cfold, Boltz-1, and Chai-1. The dataset spans diverse functional classes, including enzymes, receptors, and transport proteins (Table 1).

**Table 1.**
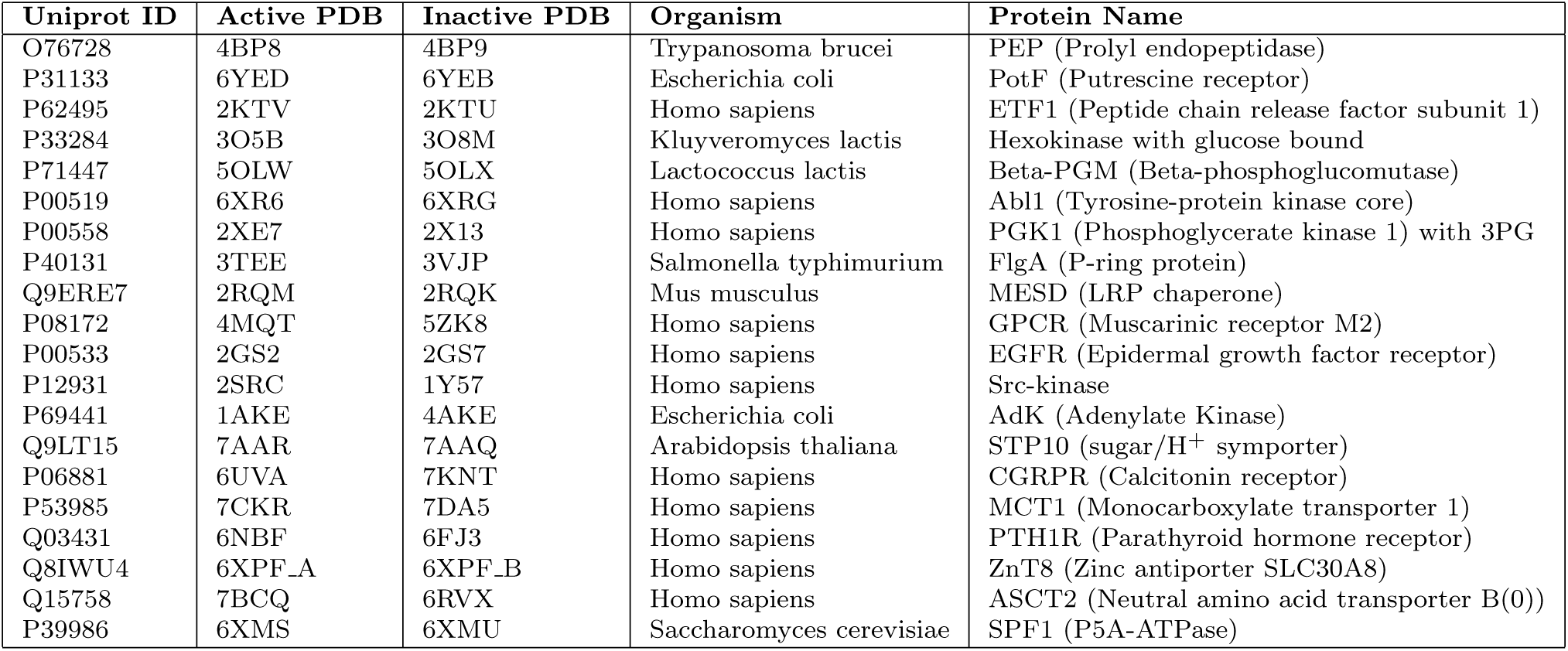
Summary of the curated protein dataset used for benchmarking for targets.

We followed Jing et al. [26] and defined precision as the average TM-score from each prediction to its closest ground truth structure, and recall as the average TM-score from each ground truth structure to its closest prediction. To investigate how well each method is able to predict the stable states across all protein targets, the average precision was plotted as a function of average recall (Fig. 1a). Chai-1 stands out as the best sampling method, achieving close to 90% average recall and precision across all targets. We next examined whether models could capture state-dependent residue-level variation by computing the Spearman correlation between per-residue C*α* RMSF in the predictions and C*α* RMSD between the active and inactive states. Chai-1 performed best, with an average Spearman correlation of 0.60 (Supplementary Fig. 1).

**Fig. 1.**
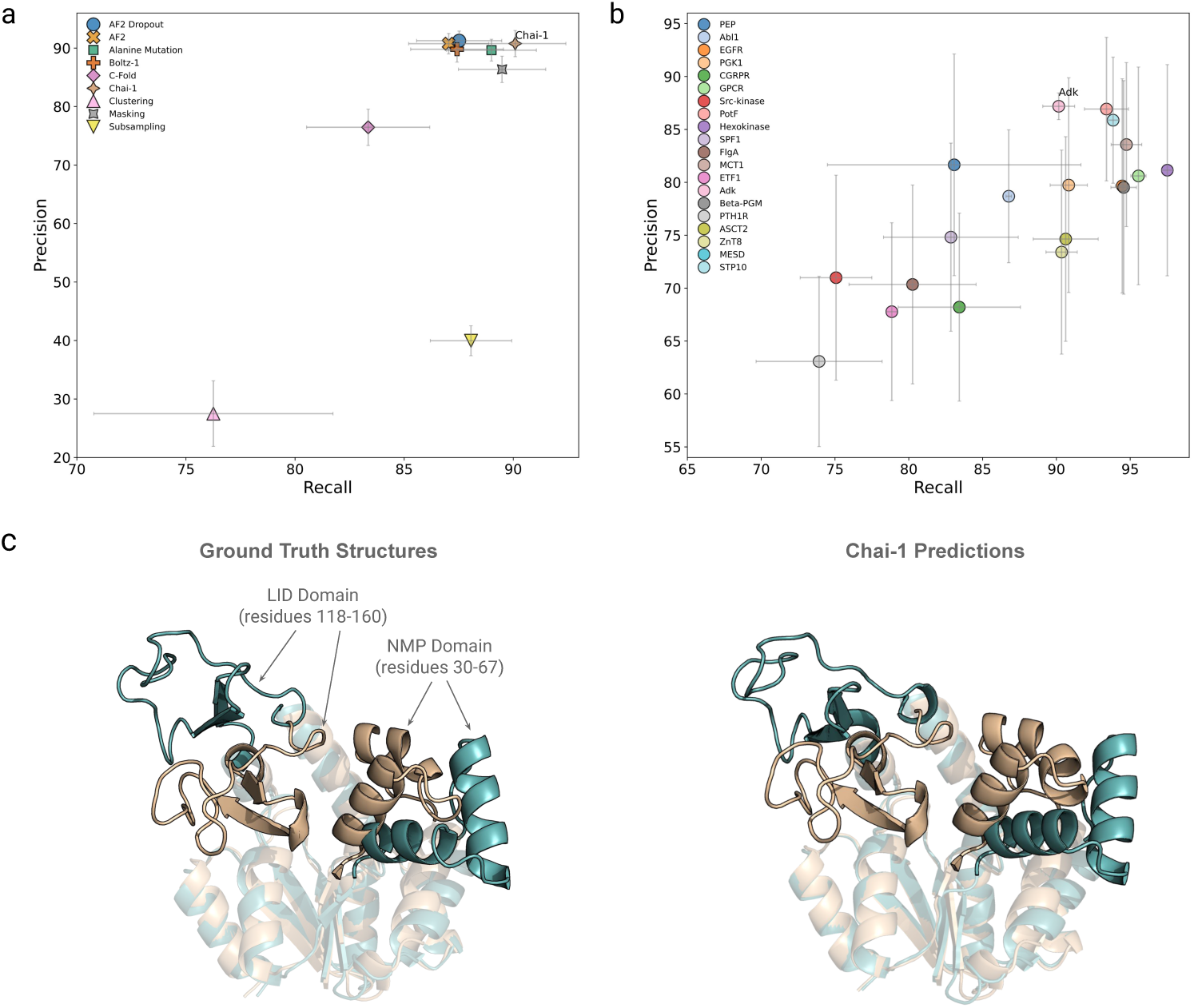
**a:** Average precision vs. recall across all targets per sampling method. Values are computed as a percentage. Standard error bars across targets are annotated. **b:** Average precision vs. recall across all sampling methods per target. Adenylate Kinase is labeled on the plot. Values are computed as a percentage. Standard error bars across sampling methods are annotated. **c:** Alignment of Adenylate Kinase (P69441) active state (orange) to inactive state (blue). The left panel shows the ground truth states (1AKE for active and 4AKE for inactive) whilst the right panel shows the closest predictions to each respective state from Chai-1 based on TM-score. Functional regions are labeled, which correspond to the predicted mobile domains.

By averaging precision and recall per protein across methods, we observed clear differences across targets (Fig. 1b), indicating that intrinsic target properties influence method performance. AdK was consistently well recovered among methods, in agreement with the residue-level analysis, where the Spearman correlation between predicted C*α* RMSF and inter-state C*α* RMSD reached 0.77 Å (Supplementary Fig. 1). Fig. 1c illustrates Chai-1’s predictions compared to the open and closed AdK functional states, demonstrating high fidelity in predictions, with a backbone RMSD of 0.61 Å and 1.59 Å to the active and inactive state, respectively. We additionally fitted a linear model with protein and method as fixed effects, and performed a two-way ANOVA. Protein explained 47.8% of the variance and method explained 13.2% (*p* < 0.001), indicating that intrinsic protein properties have a greater impact on performance than the choice of sampling method, though substantial variability remains. Residual diagnostics indicated approximate normality, homoscedasticity, and independence of residuals. Three categories of features were derived to investigate further the impact of intrinsic protein properties ( Supplementary Table 1). The relationships are visualized in Supplementary Fig. 2, with corresponding Spearman correlation values reported in Supplementary Table 1. Particularly, structural heterogeneity features (such as global and mean RMSD, average AE, and pLDDT) and sequence composition features (such as molecular weight, protein length, cysteine count, and *β*-sheet prevalence), together with MSA quality, exert a stronger influence on sampling performance than the sampling protocol itself.

### 2.2 Chai accurately recovers both active and inactive states of AdK

We selected AdK to further evaluate the accuracy of sampled conformations because it was the best-performing target across all methods and is a well-characterized model system with extensive experimental data, a rich literature in conformational dynamics, and publicly available simulation datasets. Among the tested methods, Chai-1 achieved the best balance between precision and recall (Fig. 2a), with the highest recall of 95%. However, it is worth noting that all methods perform quite well on AdK, given the low range of precision and recall values (around 85% to 100% for recall and around 80% to 95% for precision). Clustering has a high recall but relatively low precision, indicating that it is able to model both states but predicts many states in between that stray far from the ground truths, indicating noisy predictions. Cfold on the other hand has a high precision but relatively low recall, indicating that it predicts one state very accurately but misses the other state. Alanine mutation has the lowest precision and recall, indicating the method produces predictions that are structurally unrelated to both states.

**Fig. 2.**
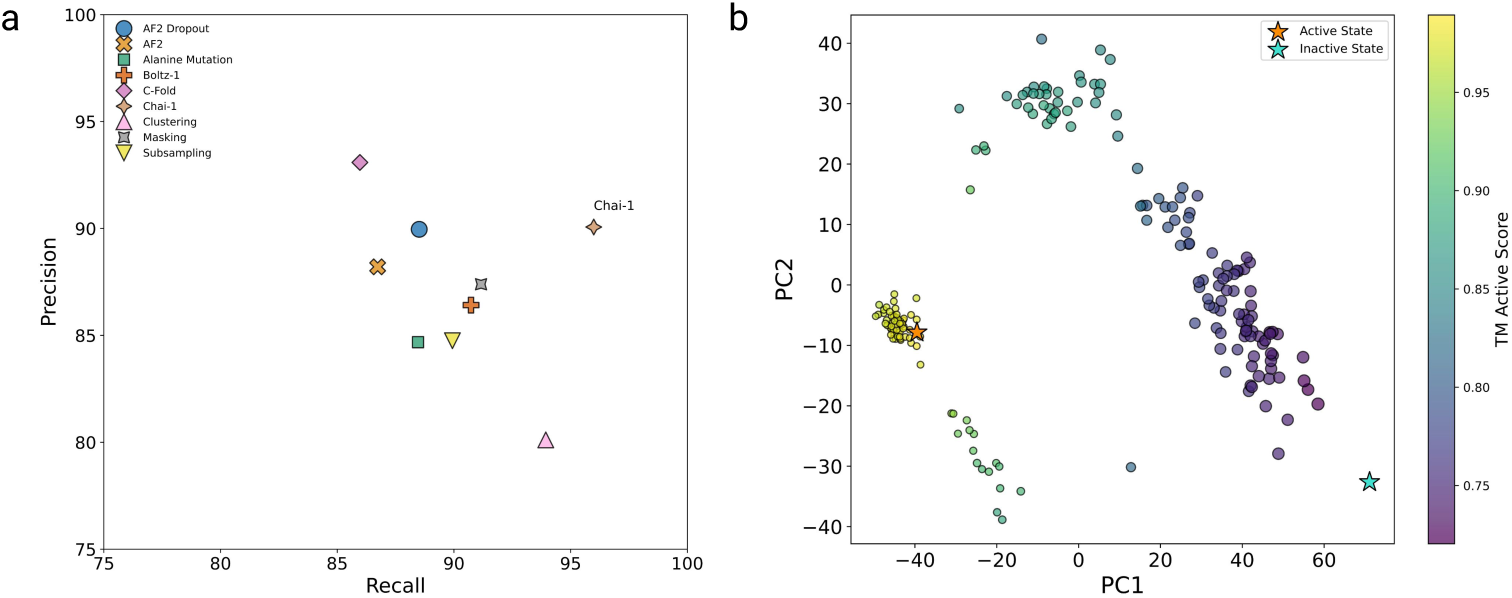
**a:** Precision vs. recall across predictions of per sampling method for Adenylate Kinase. Values are computed as a percentage. **b:** PCA plot of predictions from Chai-1. PCA was run on the C*α* xyz coordinates of all residues in each prediction, then PC2 was plotted against PC1. Each point represents one prediction, colored by the TM-score to the active state, where point size is proportional to the TM-score to the inactive state. The orange star represents the active conformation ground truth and the teal star represents the inactive conformation ground truth.

Overall, most methods consistently generate structurally accurate conformations across different sample sizes, reflected in the high precision (Supplementary Fig. 3b). In contrast, increasing sample size helps enhance coverage of the conformational landscape, reflected by the increasing recall (Supplementary Fig. 3a), highlighting that larger sample size enhances coverage of the conformational landscape rather than per-state fidelity. Chai-1 maintained high precision and recall even when the sample size was reduced. Although MSA clustering was the only method that did not plateau in recall, its scalability was limited by MSA depth.

Given that the ground truth structures for all functionally relevant conformational states are often unknown, and that some proteins may adopt more than two stable states, we performed principal component analysis on C*α* coordinates of Chai-1’s predictions on AdK and applied unsupervised clustering to study the conformation space. Distinct clusters emerged, suggesting different conformational states, in the absence of labeling or prior structural knowledge (Fig. 2b). By projecting the experimentally determined structures into the same PCA space, the active and inactive states were separated, corresponding to two distinct clusters. These results demonstrate that unsupervised methods can meaningfully capture the underlying conformational landscape, potentially enabling the identification of metastable or transitional states that are not experimentally resolved.

### 2.3 Validation of predicted intermediate conformations using molecular dynamics

As shown in Fig. 3a, Chai-1 not only captured both active and inactive conformational states but also generated intermediates that may represent transitions between them. To assess these predictions, we compared them against established transition modeling methods: Dynamic Importance Sampling MD (DIMS), which models physically realistic transitions with atomic interactions, and Framework Rigidity Optimized Dynamics Algorithm (FRODA), which uses geometric constraints to generate stereochemically plausible pathways [27]. From publicly available AdK closed-to-open transition datasets, 19,691 conformations were extracted from DIMS trajectories and 28,282 from FRODA trajectories [28]. When projected into TM-score space (Fig. 3b), both MD approaches spanned the conformational landscape between active and inactive states, with DIMS more closely following stable states and FRODA deviating slightly but retaining local minima near the inactive state and mid-trajectory.

**Fig. 3.**
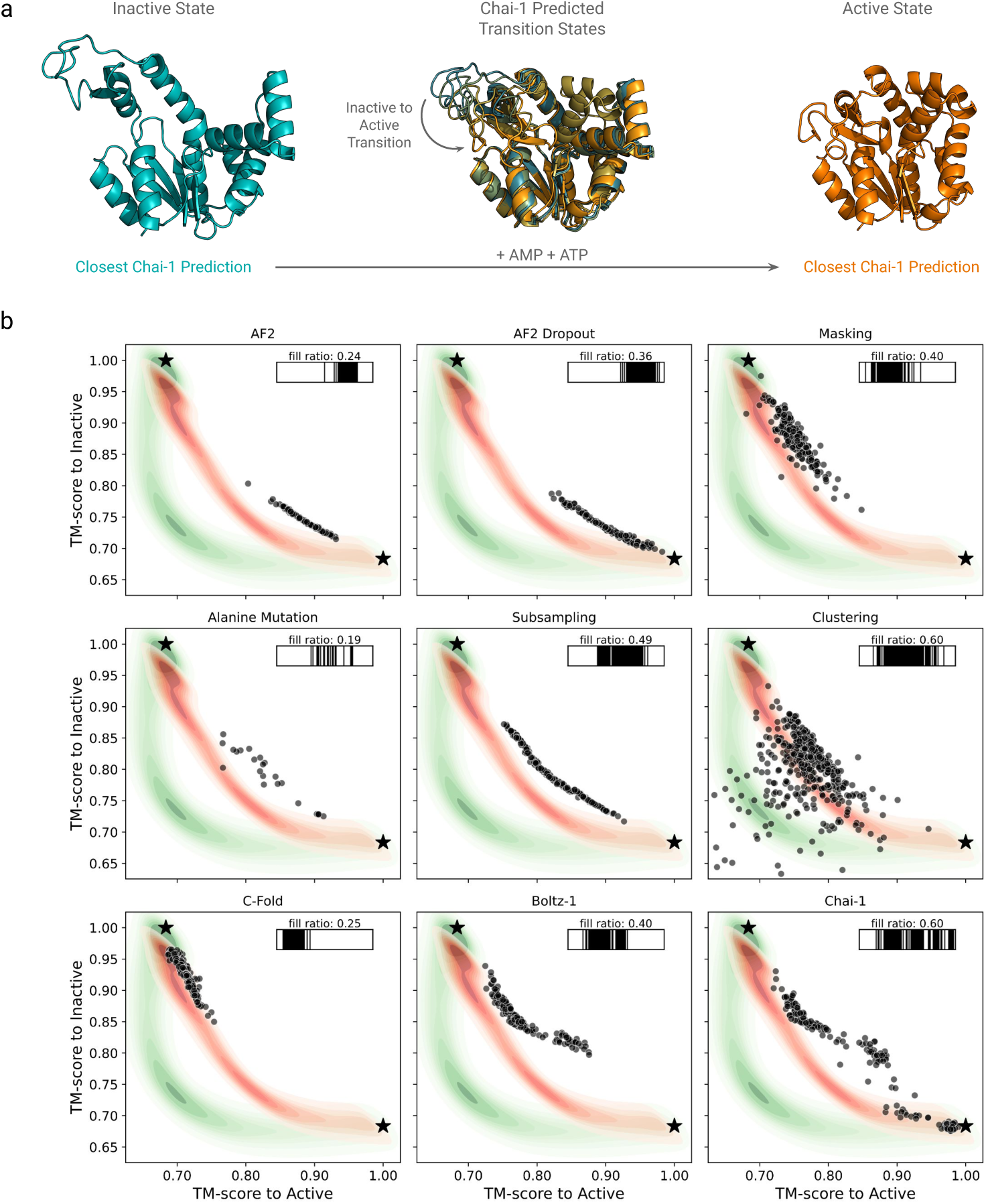
**a:** Visualization of conformational landscape of Adenylate Kinase. Transition between inactive (open) and active (closed) Adenylate Kinase conformations with intermediate states extracted from Chai-1 predictions. **b:** Per-target comparison of TM-scores to the ground-truth inactive versus active states for predictions from all sampling methods, and compared to conformations extracted from MD- and geometry-based simulations. Predictions from each sampling method are shown as black circles, and the ground-truth active and inactive structures as black stars. Conformations sampled from the MD and geometry-based trajectories are represented as spatial density clouds (red: DIMS; green: FRODA). Each subplot also includes binary histogram visualizations of the fill ratio, with black regions indicating filled bins.

Among the tested methods, Chai-1 ensembles deviated least from the MD pathways and achieved the highest fill ratio of 0.60. In line with the precision-recall analysis (Fig. 2a), Chai-1 produced low-variance predictions and most closely sampled both ground truth states. Clustering reached a comparable fill ratio but often produced noisy, low-quality conformations. Subsampling, though more limited in coverage, aligned closely with DIMS and successfully captured intermediates along the transition trajectory. By contrast, other approaches, such as Cfold and alanine mutation, generated only partial coverage, typically confined to one state or sparse intermediates, without spanning the full transition.

Projection of the C*α* coordinates into PCA space shows that the transition pathways sampled by DIMS and FRODA differ from those generated by folding model-based methods (Supplementary Fig. 4a). Specifically, Chai-1 and Subsampling trajectories diverge from DIMS, whereas DIMS and FRODA exhibit substantial overlap. The intermediate samples from Chai-1 revealed a conformational landscape for AdK in which the LID domain moves gradually while the NMP domain undergoes a delayed, abrupt movement (Supplementary Fig. 4b). This differs from the Subsampling and MD trajectories, where the two domains move more concertedly, though Subsampling fails to closely approach either end state. Notably, NMR and computational studies indicate that NMP opening typically precedes full LID motion [29–31], suggesting that the highly synchronous pathways observed in Subsampling and MD may represent less probable but mechanically possible routes within the enzyme’s dynamic landscape.

An experimentally determined intermediate structure with a semi-open LID domain was used to validate Chai-1 predictions of intermediate states (PDB ID: 1ZIN). Backbone RMSDs between this ground truth and all Chai-1 predictions reveal that the three closest predictions (RMSD < 2Å) belong to a cluster of structures centered around PC1 = –25 and PC2 = –20 in the PCA plot (Supplementary Fig. 4a, Supplementary Fig. 5). This result further supports the conclusion that Chai-1 can accurately capture intermediate conformations.

### 2.4 Conformation variation allows prediction of mobile domains

Beyond global conformations, we examined residue-level dynamics using C*α* RMSF of the predicted ensembles to evaluate whether Chai-1 captures functional domain variability, as reflected by the C*α* RMSD between AdK’s active and inactive states (Fig. 4a). The regions showing the largest divergence were highly consistent between the RMSD and RMSF profiles and corresponded to AdK’s functional domains, the NMP and LID domains. This concordance indicates that Chai-1 not only reproduces the conformational endpoints but also accurately identifies the dynamic regions of the protein, enabling recognition of mobile domains directly from the predicted ensemble. The aligned error matrix is commonly used to visualize inter-domain dynamics, with the (*i, j*)-th element defined as the RMSD of residue *j* when the active and inactive structures are aligned on residue *i*. We extended this concept to sampled conformations, by computing the same quantity across all possible pairs of samples and averaging the resulting values (Fig. 4b). Distinct off-diagonal blocks with higher RMSD values (∼3.5 Å) between residues 30-60 and 120-160 indicate coordinated displacement of these mobile domains, while low diagonal values confirm internal structural integrity.

**Fig. 4.**
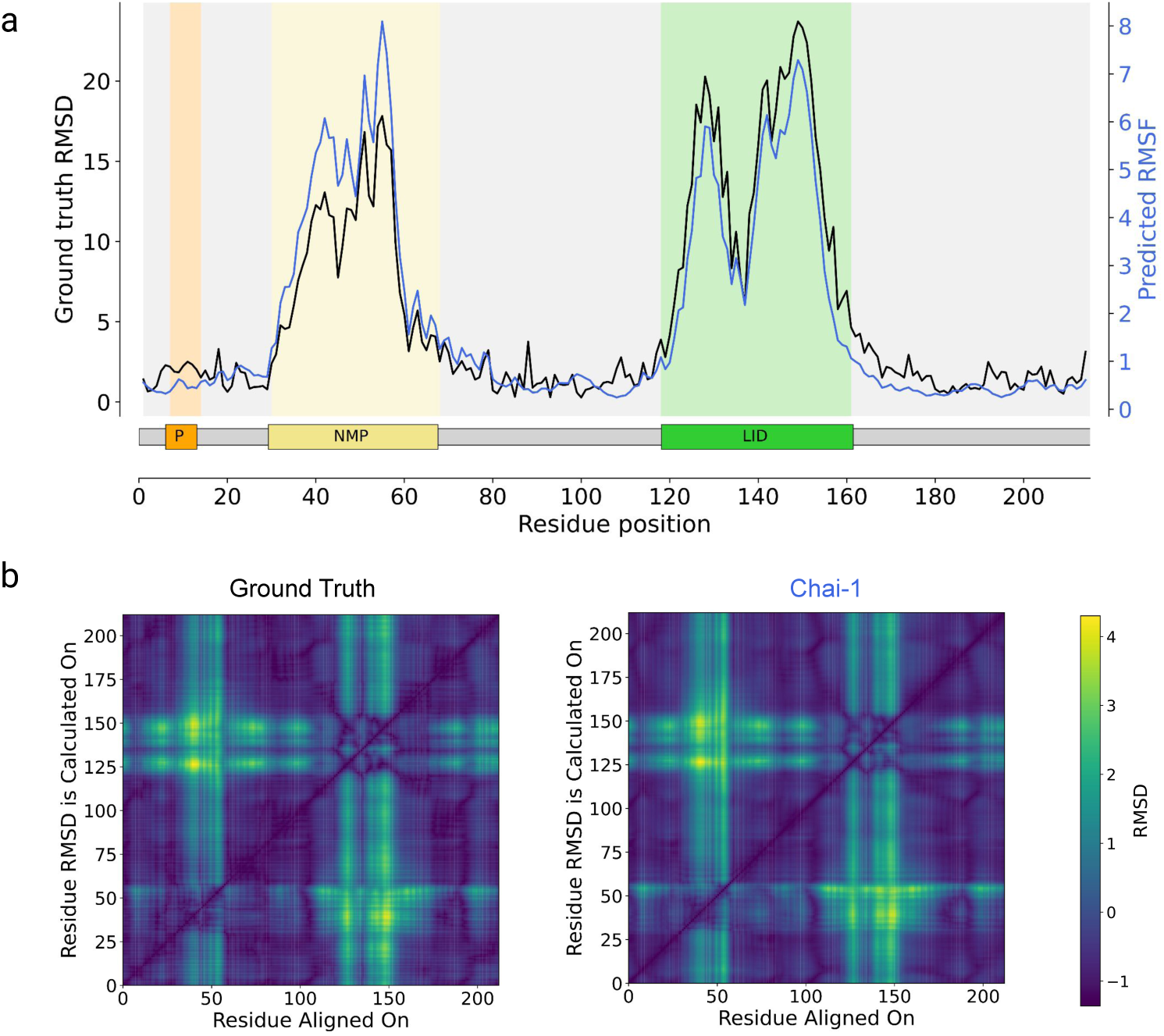
**a:** Per-residue C*α* RMSD (Å) between ground truth states plotted with per-residue C*α* RMSF (Å) across predictions from Chai-1 mode for target Adenylate Kinase (P69441). The functional regions of Adenylate Kinase are annotated, where orange represents the P-loop which is highly conserved and is involved in hydrogen bonding and cation-*π* interaction with residues in CORE and LID for stabilisation of the closed domain, the yellow represents the NMP domain which is responsible for AMP-binding, and green represents the LID domain which is responsible for ATP-binding [29]. **b:** Alignment error matrices for target Adenylate Kinase (P69441) from the ground truth states and for predictions from Chai-1. Green represents high variability in residue positions between states and across predictions, for the ground truths and predictions from Chai-1.

This pattern is observed in both the aligned error matrix and extended matrix, suggesting an accurate prediction of hinge-like motion between stable domains rather than local unfolding or disordering, consistent with prior reports [32]. The proposed method thus offers a concise view of inter-domain mobility and intra-domain stability. Supplementary Fig. 6 shows AE matrices across all sampling methods, with most performing comparably to Chai-1.

Further analysis reveals that Chai-1 is able to capture subtle mechanistic details on a residue level that are important for AdK function. Among residues of the LID domain, the lysine in position 136 was found to form an essential salt bridge interaction with the aspartic acid in position 118, largely responsible for stabilizing the open state of the protein [33]. The relationship between the D118-K136 distance and the TM-score of Chai-1 predictions relative to the active state (Supplementary Fig. 7) reveals that the distance increases as the predicted conformation approaches the active (closed) state, consistent with prior atomistic studies.

## 3 Discussion

In this work, we conducted the most extensive benchmark to date across twenty proteins and nine methods. We showed that Chai-1 successfully samples biologically relevant conformations across different proteins, outperforming other folding model-based sampling methods. Using Adenylate Kinase, a prototypical enzyme that exhibits large motions in its NMP and LID domains [29], as an example, Chai-1 not only recovered both open and closed states but also generated plausible intermediates spanning the transition. By generalizing the aligned error concept from two structures to an ensemble, we further pinpointed the D118-K136 salt bridge as a key stabilizer of the open (inactive) state.

Intermediate samples showed that Chai-1 effectively explored Adenylate Kinase’s conformational space and predicted a pathway where the LID domain moves gradually and the NMP domain undergoes a delayed, higher-barrier transition. This sequence of motions is consistent with NMR and computational studies showing that full LID opening requires the NMP domain to adopt the open conformation [29–31]. In contrast, molecular dynamics trajectories from DIMS and FRODA reported more concerted domain movements, potentially biased by simulation forcefields and constraints. Together, these results indicate that Chai-1 captures the biologically favored order of conformational change and provides a computationally efficient alternative to molecular dynamics for probing mechanistically accurate protein transitions.

During benchmarking, we discovered that intrinsic protein properties exert a stronger influence on sampling accuracy than the choice of method, an effect not previously well recognized. For example, proteins with large conformational heterogeneity, reflected by high RMSD and average aligned error between states, were typically recoverable in one state but limited in the ensemble’s coverage of both functional states. Sequence features also shaped flexibility accuracy where longer proteins, higher molecular weight, greater cysteine content, and lower *β*-sheet prevalence all reduced the ability of models to capture residue-level dynamics, indicating that protein size and composition influence predictive performance. We also found that deep and well-aligned MSAs, containing rich evolutionary information, improved recovery of both global conformational states and residue-level flexibility. These insights not only highlight previously overlooked determinants of sampling success but may also guide the selection of target proteins for conformational studies and inform strategies to improve prediction quality.

Nevertheless, important limitations remain. Current folding models are computationally expensive, and extensive conformational sampling further increases costs, motivating the development of more efficient strategies such as reusing previously generated samples or modifying the diffusion process to accelerate sampling and improve quality. The predicted ensembles do not explicitly capture the physics of the system, and incorporating force fields or other physical constraints could improve their thermodynamic realism. In addition, these ensembles lack an inherent concept of time or ordering, unlike molecular dynamics simulations; one possible future direction is to train diffusion models to generate temporally ordered pathways that transition from one conformation to another. Our analysis of domain motion was limited to a single case study protein and to monomeric targets; future work should expand to multiple targets and consider multi-chain proteins, including how one protein’s conformation may vary in the presence of another protein, such as a chaperone or binding partner. Accounting for allosteric signals, such as ions or chemical substrates, could further improve modeling of conformationally dynamic proteins. Finally, combining simulations with experimental measurements would provide complementary validation and guidance for model development.

In conclusion, we performed a rigorous benchmark of modern structure prediction models across a diverse panel of proteins and introduced a systematic analysis framework spanning the state, domain, and residue levels. Using Adenylate Kinase as a case study, we showed that Chai-1 accurately recovered both functional states and plausible intermediates. By leveraging residue-level analyses, we further showed how these predictions can be dissected to reveal mechanistically relevant motions, distinguishing global domain shifts, local rearrangements, and disorder-order transitions. These results address a gap in conformational landscape studies, particularly for diffusion-based models. More broadly, our work highlights the need to investigate conformational dynamics, often missed by static models, for understanding the mechanisms of allosterically regulated proteins. Such approaches also hold promise for translational applications, including vaccine design, where antigens can be engineered to reduce conformational variability [34].

## 4 Methods

### 4.1 Dataset curation

A set of 20 monomeric proteins with known active and inactive conformational states was curated from previously published benchmarks, prioritizing proteins used across multiple studies (see Table 1). The protein structures were preprocessed by removing heteroatoms and peptides, trimming residue ranges, and reindexing residues to ensure both conformational states had matching sequence and residue numbering.

### 4.2 MSA generation

The master MSAs used for all methods were pre-generated through MMseqs2 [35] and HHsearch [36] using the ColabFold v1.5.5 notebook [37].

### 4.3 Conformational sampling methods

All benchmarked folding model-based conformational sampling strategies were run with the same sequence and master MSA. Standard AlphaFold2 inference [11] used the model 2 ptm checkpoint with relaxation off, generating 200 predictions from different random seeds, with the same settings applied across all AlphaFold2-based methods. AlphaFold2-Dropout [24] enabled dropout in the Evoformer and structure modules at 10–25% depending on layer, producing 200 samples from different seeds. AFSample2 [22] employed a 15% MSA column masking fraction to generate 200 samples using the parameters from the original paper. SPEACH-AF [23] followed the published protocol, generating 15 predictions per sliding mutation window, with total conformations depending on sequence length. Subsampled AlphaFold2 [20] used maximum extra MSA sequences and clusters set to 16, sampling three max seq:extra seq settings (32:16, 128:64, 5120:512) to produce 200 samples per target (67 from 32:16 and 128:64, 66 from 5120:512). AF-Cluster [21] applied DBSCAN to cluster MSAs, with prediction count depending on clusters. Cfold [25] followed the original parameters to produce 200 samples. Finally, AlphaFold3-style models Boltz-1 [17] and Chai-1 [15] performed standard MSA-based inference with default parameters, generating 200 diffusion samples per target.

### 4.4 Evaluation metrics

#### 4.4.1 Precision and recall

Similar to Jing et al. [26], we calculated precision and recall for evaluating predicted ensembles. Precision is defined as the average similarity (TM-score) from each prediction to its closest ground truth structure; recall is the average similarity from each ground truth structure to its closest prediction. Compared to Jing et al. [26], we replaced lDDT-C*α* by TM-score as TM-score better captures global topology, handles domain motions (e.g., hinge movements), and penalizes incorrect domain arrangements. While lDDT can, in principle, be normalized, it remains a local metric and is less suited for assessing global conformational shifts. TM-score, by contrast, is length-normalized by design, facilitating more consistent comparisons across proteins of varying sizes.

#### 4.4.2 Structural similarity and flexibility

TM-score and root-mean-square deviation (RMSD) were used as general structural similarity metrics. To quantify residue-level flexibility across prediction ensembles, we used root-mean-square fluctuation (RMSF) values after aligning all predictions to a reference structure. Per-residue RMSD values between aligned ground truth states were also computed to characterize biologically relevant conformational shifts.

#### 4.4.3 Alignment error matrices and correlation with ground truths

Aligned error (AE) matrices were computed by aligning structures on residue *i* and calculating the RMSD of all residues *j* = 1 … *n*, where *n* is the total number of residues; diagonal elements satisfy RMSD*_ii_* = 0. For predicted structure ensembles, each pair of predictions was aligned and the same procedure was applied, producing one AE matrix per pair. These matrices were averaged over all pairs to yield a single AE matrix per protein per method. Unresolved or non-common residues between experimental and predicted structures were filtered prior to analysis. Flattened AE matrices were compared to the ground truth using Spearman correlation for each target. These correlation coefficients were then used in statistical analyses to assess the effects of the protein target and sampling method on performance.

#### 4.4.4 Fill-ratio of conformational space

The fill-ratio [22] was calculated by plotting TM-scores to both active and inactive states for all predictions and projecting each point onto the segment connecting the two ground truth states. This segment was divided into 100 bins, and the fill-ratio was defined as the fraction of bins containing at least one projected point.

## Declarations

### Data availability

The structural data that supports the findings of this benchmark study are publicly available to download from the RCSB Protein Data Bank, with the PDB IDs associated with the active and inactive states of all targets supplied in Table 1. The MSA data used in this study can be generated with the ColabFold notebook from Kim et al. [37]. The DIMS and FRODA trajectory data for Adk from Beckstein et al. [28] are available in figshare.

### Code availability

In this study, the AlphaFold2-based methods were implemented using OpenFold, and the code is available at https://github.com/instadeepai/FoldConfBench. For Chai-1 and Boltz-1, the corresponding open-sourced code was used and run with an A100 NVIDIA GPU with 40GB memory. The default model parameters were used and 200 samples were generated per AlphaFold3-based method. The source code for Chai-1 and Boltz-1 can be found at https://github.com/chaidiscovery/chai-lab and https://github.com/jwohlwend/boltz, respectively.

## Supplementary Figures & Tables

**Fig. S1.**
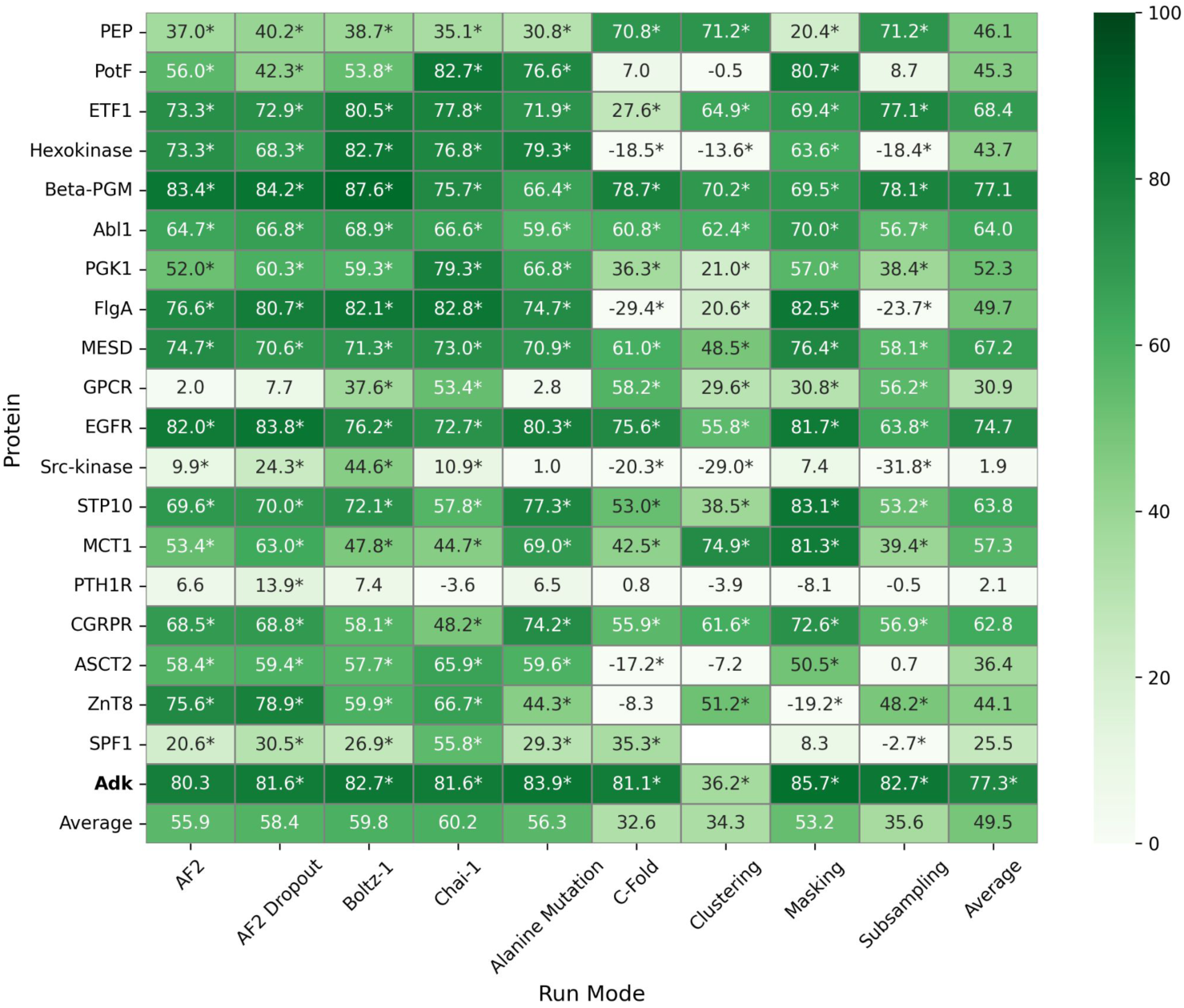
Heat map summarizing Spearman correlations between per-residue ground-truth C*α* RMSD (Å) and C*α* RMSF (Å) across predictions for each target and sampling method. Asterisks (*) indicate statistical significance at p < 0.05. No value was obtained for P39986 from clustering due to only a single cluster being returned.

**Fig. S2.**
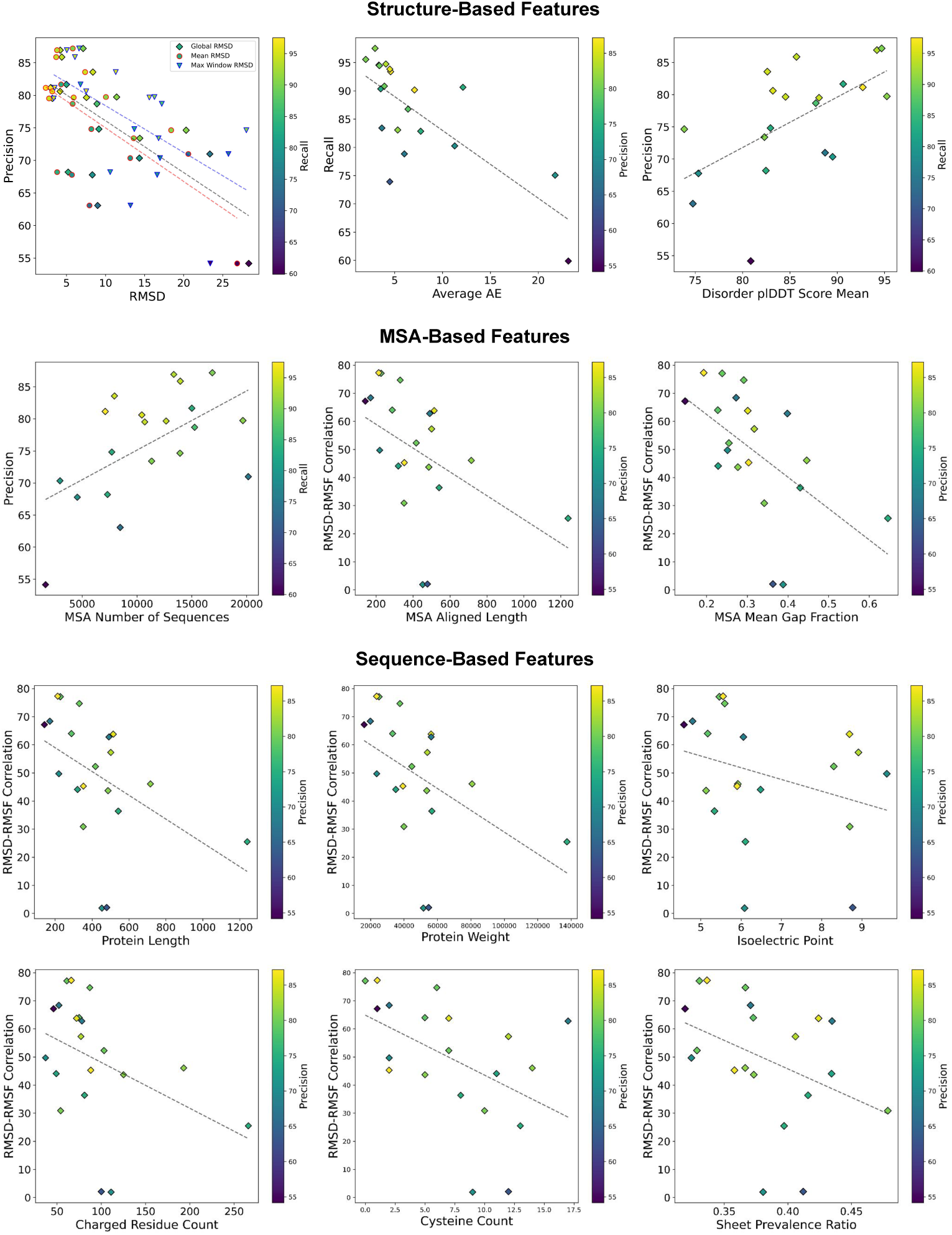
Correlation between target-based features and performance metrics. Plots show correlations of selected features across all three categories with each primary metric (precision, recall, or RMSD–RMSF correlation; y-axis). In each plot, points are additionally colored by a different label (as indicated by the color bar).

**Fig. S3.**
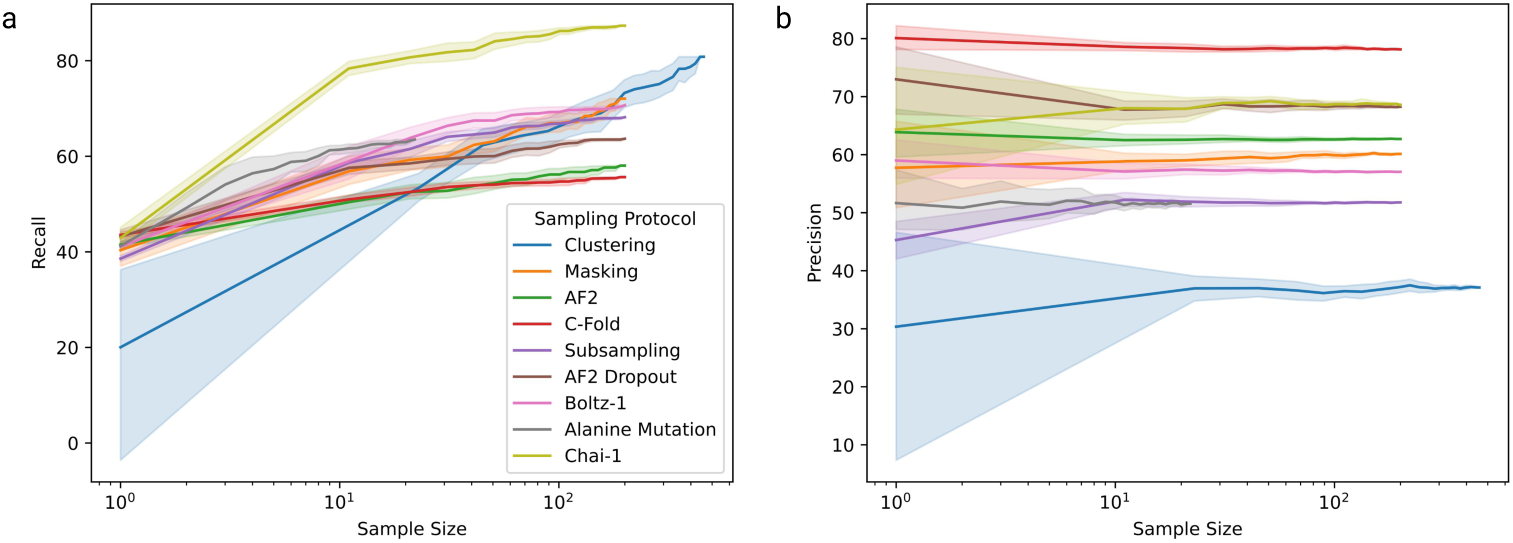
**a:** Average Recall vs Sample Size across adenylate kinase predictions for all sampling protocols. **b:** Average Precision vs Sample Size across adenylate kinase predictions for all sampling protocols.

**Fig. S4.**
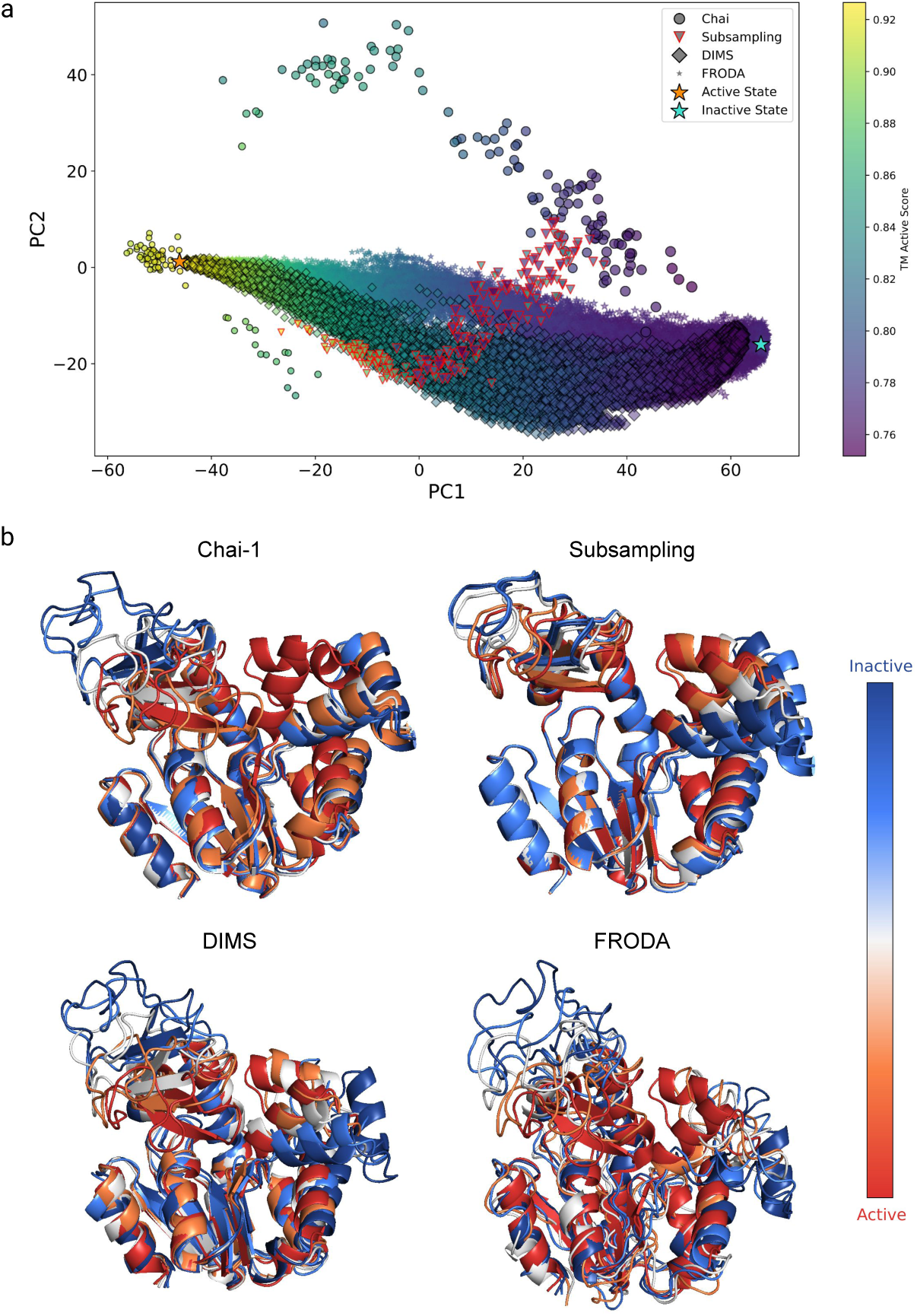
**a:** PCA plot of predictions from Chai-1 and Subsampling with extracted frames from DIMS and FRODA. PCA was run on the C*α* xyz coordinates of all residues from the predictions from Chai-1 and Subsampling with the ground truth states. Then the C*α* xyz coordinates of all residues from the DIMS and FRODA frames were projected into the same space. PC2 was then plotted against PC1. Each point represents one conformation, colored by the TM-score to the active state, where point size is proportional to the TM-score to the inactive state. The orange star represents the active conformation ground truth and the teal star represents the inactive conformation ground truth. **b:** Conformations along PC1 colored from active (red) to inactive (blue). Five conformations were extracted along PC1: those closest to the active and inactive states at the extremes, and three intermediates evenly spaced between them.

**Fig. S5.**
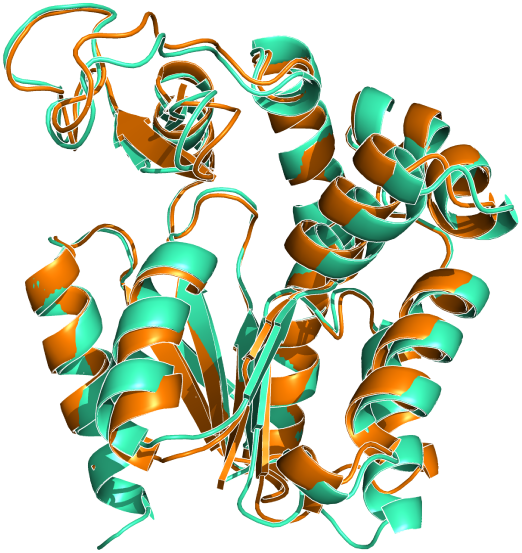
Alignment of an experimentally-resolved Adk intermediate structure (PDB: 1ZIN, green) and the closest Chai-1 predicted intermediate state (orange), backbone RMSD = 1.94Å.

**Fig. S6.**
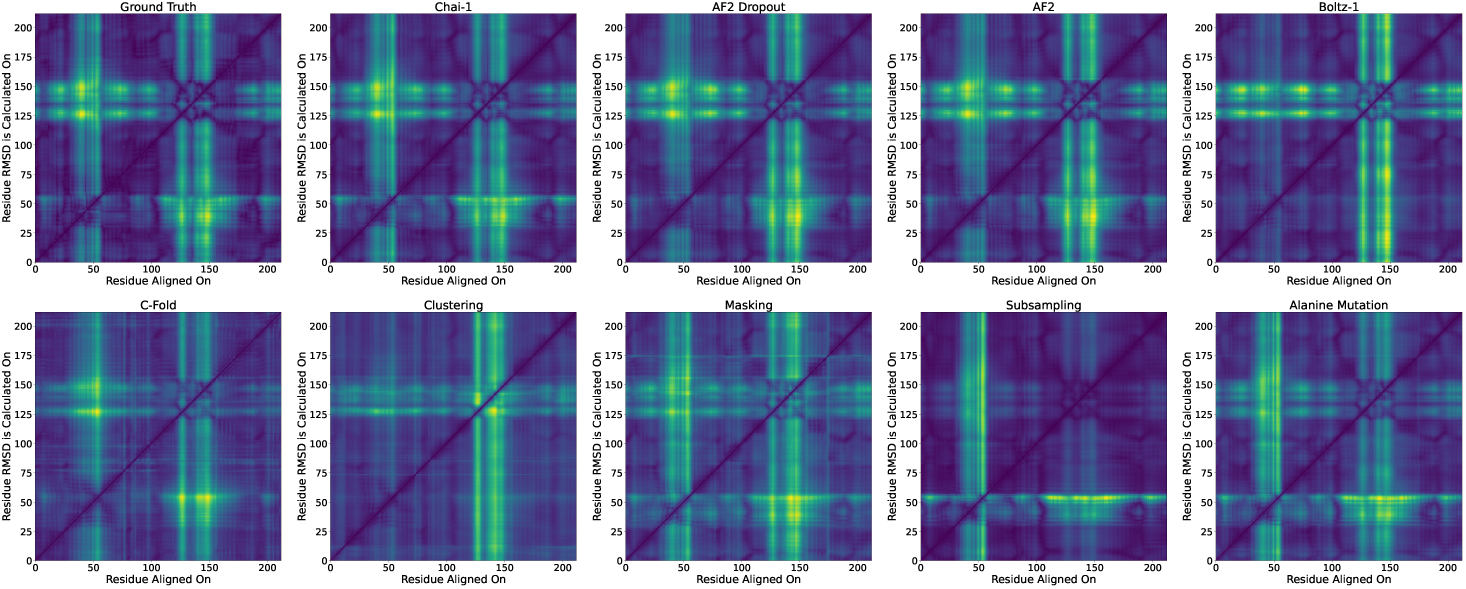
Alignment error matrices for target adenylate kinase (P69441) from the ground truth states and for predictions from all sampling protocols. Green represents high variability in residue positions between states and across predictions.

**Fig. S7.**
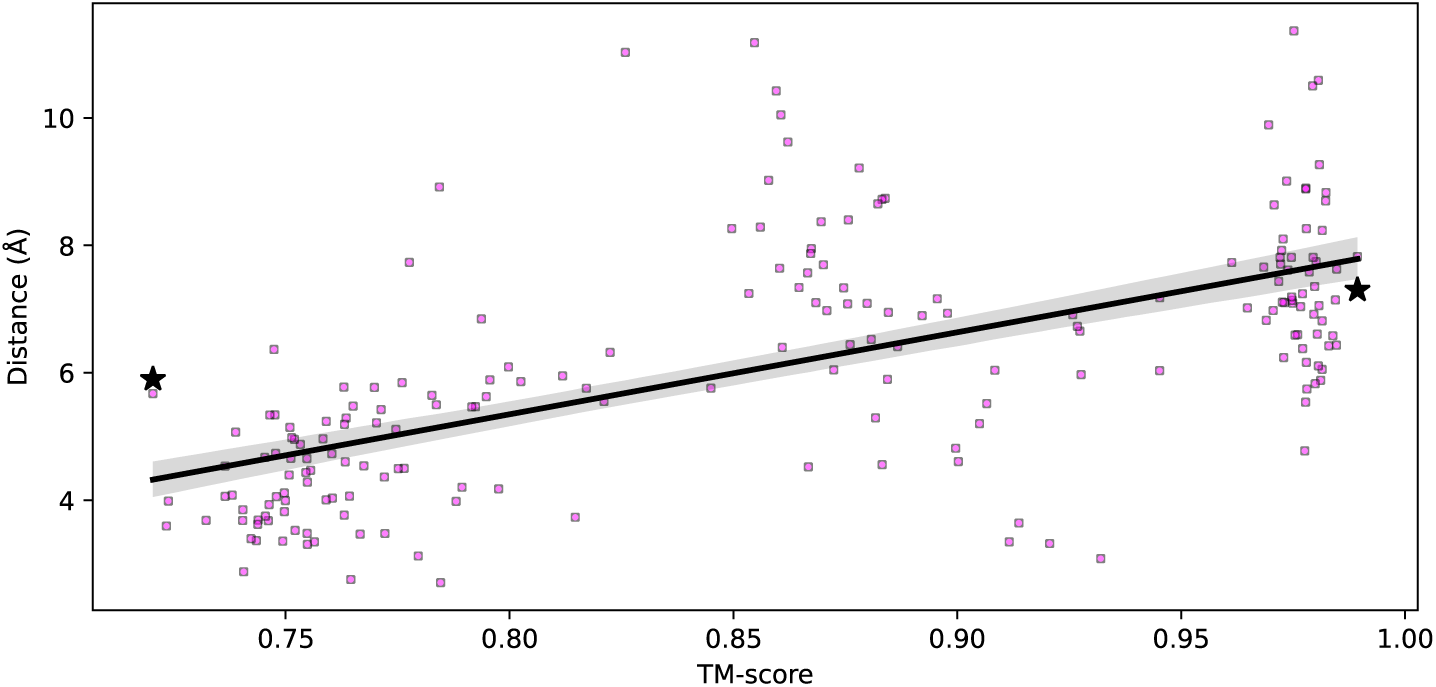
Relationship between the distance of the D118–K136 salt bridge and the TM-score of Chai-1 predictions relative to the active state.

**Table S1.** Spearman correlations between target-based features and performance metrics (precision, recall, and RMSD–RMSF correlation). Asterisks (*) indicate statistical significance at p < 0.05. Target features are grouped into three categories: (1) structure-based features: global C*α* RMSD between aligned ground-truth structures, mean per-residue C*α* RMSD, maximum sliding-window C*α* RMSD (over 20 residues), average alignment error (AE), and disorder pLDDT score; (2) MSA-based features: number of sequences in the master MSA, aligned length (number of alignment positions including gaps), and mean gap fraction (average proportion of gaps across all sequences in the MSA); and (3) sequence-based features: protein length, molecular weight, isoelectric point, charged residue count, cysteine count, and *β*-sheet prevalence ratio.

| Protein Feature | Precision | Recall | RMSD-RMSF Corr. |
| --- | --- | --- | --- |
| Global RMSD | -0.61* | -0.65* | -0.24 |
| Mean RMSD | -0.52* | -0.60* | -0.24 |
| Max Window RMSD | -0.65* | -0.57* | -0.18 |
| Average AE | -0.31 | -0.73* | -0.06 |
| Disorder pLDDT Score | 0.65* | 0.30 | 0.14 |
| MSA Number of Sequences | 0.49* | 0.15 | -0.03 |
| MSA Aligned Length | 0.26 | 0.19 | -0.56* |
| MSA Mean Gap Fraction | 0.02 | -0.03 | -0.64* |
| Protein Length | 0.26 | 0.19 | -0.56* |
| Protein Weight | 0.17 | 0.12 | -0.59* |
| Isoelectric Point | 0.11 | 0.07 | -0.42* |
| Charged Residue Count | 0.26 | 0.06 | -0.48* |
| Cysteine Count | -0.08 | -0.07 | -0.59* |
| Sheet Prevalence Ratio | -0.04 | 0.22 | -0.48* |

